# Deep learning enables feature extraction of 3D collagen architecture in cleared fibrotic tissues

**DOI:** 10.64898/2026.02.25.707675

**Authors:** Wout Houbart, Laura Schelfaut, Angeliki D. Vavladeli, Noah Borges, Marthe Boelens, Claudia M. Brenis Gómez, Bram Verstappe, Mohammad Ghiasloo, Nikita Vladimirov, Phillip Blondeel, Charlotte L. Scott, Fabian F. Voigt, Bart N. Lambrecht, Fritjof Helmchen, Esther Hoste, Kris Vleminckx, Thomas Naert

## Abstract

Light-sheet fluorescence microscopy enables deep optical sectioning of large, cleared biological tissues. However, effective clearing of collagen-rich tissues remains a persistent technical challenge. Moreover, standardized workflows integrating three-dimensional imaging with computational analysis of collagen architecture are currently unavailable. Here, we present an integrated pipeline combining optimized tissue clearing, volumetric light-sheet imaging, and deep learning-based feature extraction of collagen architecture. Using experimental desmoid tumors as a proxy for collagen-dense tissues, we optimised DISCO-based clearing incorporating Fast Green FCF for collagen labelling. We achieved optical transparency and full 3D visualization of collagen architectures in desmoid tumors, human skin biopsies, and fibrotic mouse lung & liver tissues, including FFPE samples. Using the Benchtop mesoSPIM platform, we acquired high-resolution volumetric datasets and validated multimodal collagen imaging through two-photon microscopy with the Schmidt-Voigt objective. To enable automated feature extraction from these large volumetric datasets, we developed ColNet, a U-Net model for automated collagen fiber segmentation. ColNet demonstrated robust generalization across diverse human and mouse tissues without retraining or hyperparameter adjustment. This integrated workflow provides a foundation for future quantitative assessment of cell-extracellular matrix dynamics in a fibrotic context.

## INTRODUCTION

Collagens represent the most abundant protein family in the human body, comprising up to 30% of total protein content. To date, 44 genes are known to encode 28 collagen types, with fibrillar collagens (types I, II, III, V and XI) representing the largest subgroup (90%)^1^. As the predominant protein family in the extracellular matrix (ECM), collagens provide structural integrity to developing tissues and organs, and serve as important regulators of tissue homeostasis^2^. Importantly, excessive deposition of collagen is linked to a wide range of fibrotic disorders (*e*.*g*. Dupuytren’s disease, idiopathic pulmonary fibrosis, osteogenesis imperfecta amongst many others), wound healing and cancer^3–5^. In cancer, the ECM constitutes an essential part of the tumor microenvironment (TME), now recognized for being crucial in cancer progression^6,7^. Aberrantly high collagen content has been linked to ECM stiffness and tumor coldness, affecting critical processes including tumor progression, immune cell infiltration, cell signaling, metastasis, and metabolic reprogramming^8,9^. Moreover, the role of ECM topology within a TME is gaining more recognition for its association with tumor cell behavior^1,10^. As such, analysis of collagen architecture ideally employs three-dimensional (3D) approaches, as two-dimensional (2D) methods are unable to fully capture tumor-matrix dynamics, depth-dependentfiber organization or ECM remodelling patterns.

Light-sheet fluorescence microscopy (LSFM) (or selective plane illumination microscopy (SPIM)), has emerged as the ideal imaging platform for volumetric analysis of complex biological structures in 3D, enabling deep optical sectioning (>1 mm) with minimal photobleaching and phototoxicity, far exceeding the limited penetration depth (<100 μm) of conventional confocal microscopy^11,12^. From a technical perspective, LSFM of dense tissues ideally requires tissue clearing protocols that achieve optical tissue transparency through depigmentation and refractive index matching. Established methods including CLARITY, CUBIC and DISCO variants render tissues transparent while preserving their architecture^13–18^. Indeed, the integration of tissue clearing with light-sheet microscopy has demonstrated versatile applications across developmental biology, neuroscience and disease phenotyping^17,19–22^.

However, a significant challenge remains the effective clearing of tissues with excessive collagen deposition, due to inadequate refractive index matching by tightly packed collagenfibers, limited reagent penetration through the dense ECM, and increased light scattering of stromal collagen^23,24^. Among the various clearing approaches, organic solvent-based methods such as DISCO demonstrate superior performance compared to hydrophilic protocols through harsher delipidation that facilitates deeper penetration^23,25^. While modified benzyl alcohol/benzyl benzoate (BABB)^26^ and Skin-iDISCO +^27^ protocols have been successfully applied to skin and soft tissue with moderate ECM content, they have not been systematically applied to the extreme collagen density of pathological fibrosis, where collagen networks directly drive disease progression. Furthermore, clearing studies primarily focus on enabling visualization rather than feature extraction of collagen architecture.

As such, structural analysis of collagen architecture represents a second critical bo leneck. Established tools such as CT-FIRE^28^, CurveAlign^29^, OrientationJ^30^, and FibrilTool^31^ have made important contributions to collagen fiber analysis, yet they were developed before the deep learning revolution. Consequently, they still require extensive parameter tuning, lack standardized pipelines, and are predominantly limited to 2D analysis. The recent advances in deep learning, and particularly the convolutional neural network (CNN) architecture U-Net, have revolutionized biomedical image segmentation by enabling automated feature extraction with minimal training data^32^. As such, many deep learning models today employ the U-Net architecture for biomedical segmentation or classification tasks, and can even be applied to phenotyping of embryonic development and disease^21,33–35^.

Using a *Xenopus* model of aggressive fibromatosis (desmoid tumor) as our technical model system for collagen-rich tissues^36,37^, we present an integrated approach that combines robust DISCO-based tissue clearing with multimodal visualization of collagen. Leveraging state-of-the-art light-sheet microscopy^20^ and two-photon imaging using the Schmidt-Voigt objective^38^, we acquired high-resolution volumetric datasets of cleared collagen-rich tissues. Based on these datasets, we trained ColNet, a U-Net-based deep learning solution for automated 3D segmentation of collagen architecture. We demonstrate that ColNet accurately segments collagen structures across diverse tissue types and pathological states, including *Xenopus* desmoid tumors, human skin biopsies and fibrotic mouse lung & liver tissue from archived FFPE material, without requiring model retraining or hyperparameter fine-tuning. With ColNet, we provide an accessible pipeline for 3D assessment of collagen organization and tumor-matrix dynamics in fibrotic malignancies.

## RESULTS

Three-dimensional visualization of collagen architecture in aggressive fibromatosis using DISCO clearing and multimodal collagen imaging Desmoid tumors, also called aggressive fibromatosis, are locally aggressive soft tissue neoplasms, marked by excessive deposition of collagen^4^. To optimize clearing of collagen-dense samples, we employed a CRISPR/Cas9 *Xenopus* model of aggressive fibromatosis^36^, which provided substantial tissue volume with pathologically elevated collagen deposition^36^ (Fig. 1a).

**Fig. 1.**
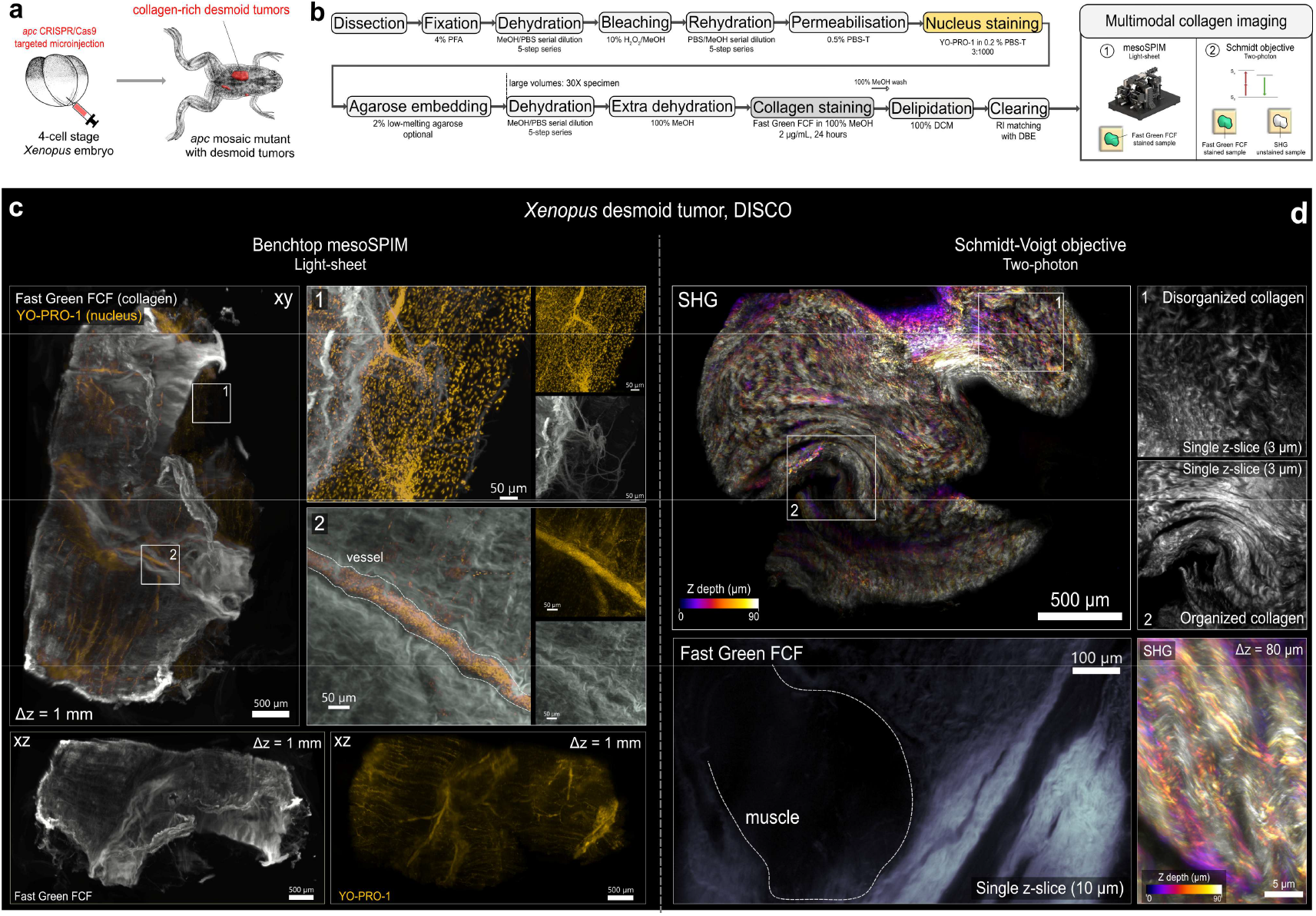
Multimodal 3D imaging of collagen architecture in a DISCO-cleared desmoid tumor.a, Schematic overview of CRISPR/Cas9-mediated desmoid tumor generation in *Xenopus*. b, Overview of the DISCO pipeline used for tissue clearing, collagen staining and visualization of desmoid tumors. Synthetic dyes Fast Green FCF and YO-PRO-1 were used for collagen and nucleus staining, respectively. Tumors were imaged with light-sheet microscopy (Benchtop mesoSPIM) or two-photon microscopy (Schmidt-Voigt objective). c, Light-sheet imaging of a cleared desmoid tumor, stained for collagen (Fast Green FCF) and nuclei (YO-PRO-1). Upper row: xy views showing maximum intensity projection (MIP) of 351 z-slices spanning 1 mm z-depth (Δz = 1 mm), with two insets displaying regions of interest at higher magnification (merged and individual channels). Bottom row: xz views of the individual Fast Green FCF and YO-PRO-1 channels. Gamma correction (1.5 in Imaris) was applied on the Fast Green FCF channel for visualization. The YO-PRO-1 16-bit greyscale map was pseudo-colored (Imaris) with a yellow look-up table (LUT) (Imaris) d, Two-photon imaging of cleared desmoid tumors using the Schmidt objective. Upper row: Depth-coded MIP of a large unlabeled desmoid tumor imaged via second harmonic generation (SHG) (925 nm excitation) spanning 90 µm. Adjacent panels (insets 1,2) show representative single z-slices (10 µm thickness) from two regions of interest (ROIs) illustrating heterogeneous collagen organization (disorganized versus organized). Lower row, left: Single z-slice (10 µm) of Fast Green FCF-stained collagen (900 nm excitation). The original 16-bit grayscale map was pseudo-colored with Bone LUT (Imaris). Gamma correction (Imaris) of 1.5 was applied on the Fast Green FCF channel for visualization. Lower row, right: Depth-coded MIP of high-magnification SHG imaging (925 nm excitation) resolving individual collagenfibers spanning 80 µm depth. All samples were imaged in dibenzylether for refractive index matching. Stitching was performed with BigStitcher. PFA: paraformaldehyde, MeOH: methanol, PBS-T: PBS with Triton-X-100, RI: refractive index, SHG: second harmonic generation.

For clearing of desmoid tumor samples, we modified the existing DISCO clearing protocol from Timin & Milinkovitch, incorporating dual fluorescent labeling with Fast Green FCF and YO-PRO-1 for collagen and nuclei labelling, respectively (Fig. 1b, Fig. S1a)^39,40^. Fast Green FCF staining was performed in 100% methanol, which was previously shown to ensure high specificity for collagen in dehydrated conditions^39,40^. We observed that limiting staining duration to 24 hours prevented oversaturation of labelled collagen while maintaining adequate signal intensity. Additionally, continuous rotation during incubation was crucial to ensure uniform dye penetration throughout the sample. Critical to successful clearing was extended dehydration in large volumes (>30X specimen volume) prior to delipidation with dichloromethane (DCM). Insufficient dehydration, whether from inadequate methanol washes or small working volumes, resulted in incomplete clearing and tissue opacity, making samples unsuitable for volumetric imaging. Following this optimized approach, we successfully cleared desmoid tumor tissue and employed both one-photon and two-photon imaging systems for visualization of collagen architecture (Fig. 1b).

Light-sheet fluorescence imaging on the Benchtop mesoSPIM^41^ enabled deep tissue penetration (up to 1 mm in our sample) and allowed, for thefirst time to our knowledge, three-dimensional (3D) visualization of collagen architecture in a desmoid tumor at cellular resolution (Fig. 1c). Fast Green FCF signal showed collagen labelling throughout the tumor, revealing a complex, dense collagen network. Individual collagen fibers were clearly resolved at higher magnifications, providing detailed insight into local fiber density & topology (Fig. 1c, insets 1 & 2). YO-PRO-1 nuclear staining simultaneously delineated the surrounding muscle tissue, revealing an organized architecture. Notably, YO-PRO-1 also highlighted a blood vessel within the tumor microenvironment by labelling of nucleated erythrocytes (a feature characteristic of *Xenopus*)^42^, with Fast Green FCF revealing dense collagen fibers wrapped around the vessel wall (Fig. 1c, inset 2). Indeed, this high-magnification view revealed the heterogeneous nature of collagenfiber organization within the tumor, ranging from irregular, invasive fibers at the tumor-muscle interface (Fig. 1c, inset 1) to more aligned, sheath like arrangements in the perivascular region (Fig. 1c, inset 2).

Next, we applied two-photon excitation microscopy using the recently introduced multi-immersion Schmidt-Voigt objective, which is ideally suited for high-resolution recordings of large, cleared samples^38^. As such, cleared desmoid tumors, either labelled with Fast Green FCF or unlabelled, could thus be directly transferred for two-photon imaging in dibenzylether (DBE) without any additional processing (Fig. S2b,c).

For unlabelled samples, we determined an optimal excitation wavelength of 925 nm for high-contrast second harmonic generation (SHG) imaging (Fig. 1d). This approach achieved full-depth penetration through 1.7 mm of dense collagenous tissue. SHG imaging revealed the heterogeneous collagen organization within the tumor, distinguishing regions of highly aligned collagen fibrors from areas of less organized collagen (Fig. 1d, insets 1 & 2), consistent with our light-sheet fluorescence observations. The endogenous SHG signal provided high-resolution imaging of collagen fibrors throughout the entire tissue volume without requiring exogenous labelling. Furthermore, two-photon visualization of collagen was performed at 940 nm in Fast Green FCF-stained samples. Further, subsequent excitation at 900 nm permi ed YO-PRO-1 visualization (Fig. S2). However, we observed spectral bleed-through of the SHG signal into the YO-PRO-1 channel. This should be considered when interpreting YO-PRO-1 nuclear channel data in collagen-rich regions under two-photon excitation.

In conclusion, the modified DISCO protocol, incorporating extensive dehydration, enables effective clearing of collagen-dense desmoid tumors while preserving compatibility with fluorescent labeling of nuclei and collagen. Combined with light-sheet fluorescence imaging and two-photon excitation via the Schmidt-Voigt objective, this establishes a multimodal imaging platform for high-resolution 3D visualization of desmoid tumor collagen architecture.

### ColNet enables 3D segmentation of collagen fibrors in a cleared desmoid tumor

To enable unbiased and automated extraction of collagen fibrors from volumetric light-sheet datasets, we trained ColNet, a deep learning model for 3D collagen fibror segmentation. Recently, deep learning for microscopy has become more accessible by platforms such as ZeroCostDL4Mic^43^ and DL4MicEverywhere^44^, providing accessible, user-friendly interfaces for training and deployment of deep learning models through cloud-based computing or remote GPU connection, eliminating the need for local high-performance computing resources and extensive computational expertise. We employed the 2D-U-Net architecture from the open-source ZeroCostDL4Mic Jupyter notebook database, which enabled straightforward model training and deployment.

ColNet was trained from scratch with a sparse annotation strategy, leveraging U-Net’s ability to train effective models with small datasets^21,32^. Following this approach, a small training dataset (n=10) of 2D slices, distributed across different depths of a light-sheet dataset (311 z-slices, 2 µm step) from a Fast Green FCF-stained cleared desmoid tumor, was manually annotated for training (Fig. S3a). Given the complex organisation of collagen fibrors, with fibrors oriented in multiple planes, we focused annotations on individual fibrors that were predominantly in-plane and clearly resolved. This approach prioritized training the model to recognize well-defined fibror structures while avoiding ambiguous annotations of out-of-plane or overlapping fibrors that could introduce noise into the training data. Using the same annotation strategy, a small test dataset (n=9) was annotated (Fig. S3b). Critically, this test dataset was withheld during training, and included not only unseen desmoid tumor sections (n=7) but also human full-thickness skin biopsies (n=2) with different collagen fibror topology, enabling assessment of the model’s ability to generalize across different tissue contexts. To maximize training efficiency with limited annotated data, we employed data augmentation during model training, including shifts, zoom, rotations, shear and flipping (Fig. S4). These image transformations artificially expanded biological variance in the training dataset and improved model performance^32^.

A critical consideration during model training is optimisation of parameters such as patch size and the learning rate schedule, which profoundly impact both convergence dynamics of loss values, andfinal segmentation quality. As such, we systematically evaluated model dynamics at different patch sizes to optimize model performance (Fig. S5). We observed that the largest patch size (within GPU memory constraints of a consumer-grade NVIDIA GeForce RTX 4090 GPU) of 1536 × 1536 pixels gives the most stable balance of local feature extraction and contextual information, leading to optimal convergence of training and validation loss. Next, we evaluated three different learning rate schedules, comparing training stability, quantitative performance metrics, and qualitative segmentation accuracy (Fig. S6). All three models achieved similar quantitative performance on the test dataset (Intersection over Union (IoU) range: 0.56-0.57, F1 range: 0.71-0.72). However, detailed examination of training dynamics and 3D segmentation predictions revealed substantial differences in qualitative performance. Model 1, for which training was ended at a learning rate of 1E-5, exhibited early convergence with incomplete optimization, resulting in undersegmentation. Model 2, trained to an ultra-low learning rate of 5E-6, showed training instability characterized by dramatic loss fluctuations and an overfitting event, resulting in oversegmentation. Model 3, employing afinal learning rate of 1E-5, with extended training at 2E-5, demonstrated stable convergence of training and validation loss without overfitting. These results demonstrate that quantitative overlap metrics alone can be insufficient for model selection in biological imaging applications, as they may mask important qualitative differences in segmentation. We selected Model 3 (final IoU: 0.57, F1 score: 0.72) for all downstream analyses (Fig. 2a).

**Fig. 2.**
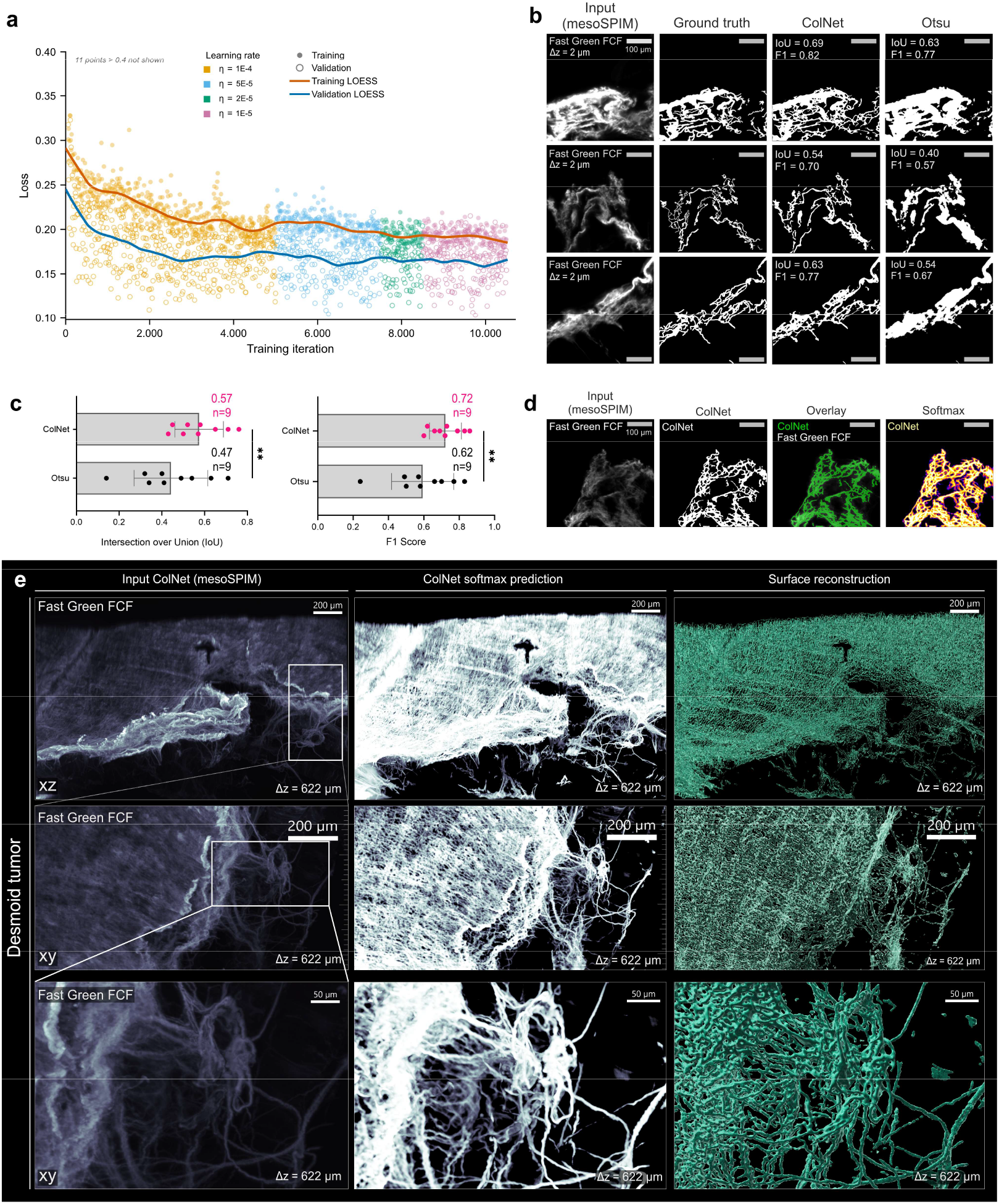
ColNet enables three-dimensional segmentation of collagen architecture in a DISCO-cleared desmoid tumor. a, ColNet training log showing training (orange) and validation (blue) loss over iterations with locally estimated sca erplot smoothing (LOESS). Loss values are color-coded for decreasing learning rates (1E-4, 5E-5, 2E-5, 1E-5). Values above 0.40 are not shown for visualization. b, Comparison of ColNet performance versus Otsu-intensity thresholding on three representative images from the training dataset. Each row shows (left to right): input image acquired on Benchtop mesoSPIM (1536 × 1536 pixels) of Fast Green FCF-stained desmoid tumor, manually annotated binary ground truth mask, ColNet segmentation mask, and Otsu segmentation mask. Intersection over Union (IoU) and F1 scores are displayed for each segmentation. c, Quantitative comparison of ColNet performance versus Otsu intensity thresholding on unseen test images (t-test, p<0.05, n=9). d, Representative example of ColNet performance on unseen test data. From left to right: input image acquired on Benchtop mesoSPIM (1536 × 1536 pixels), ColNet binary segmentation mask, overlay of input and mask, and ColNet softmax probability map. The original 16-bit grayscale softmax map was pseudo-colored with a Fire look-up table (LUT) (Fiji). e, Three-dimensional visualization of ColNet performance on desmoid tumor. Left: Maximum intensity projections (MIP) of Fast Green FCF-stained desmoid tumor acquired on Benchtop mesoSPIM, spanning 622 µm z-depth (Δz = 622 µm) Middle: ColNet softmax probability maps. Right: 3D surface renderings (Imaris) of softmax probability maps at different magnifications. Original grayscale data were pseudo-colored with Bone LUT (left and middle panels) or cyan LUT (right panel) in Imaris. Raw mesoSPIM data (left panel) were gamma-corrected (1.5 in Imaris) for visualization..

When deployed on the unseen test dataset (n=9), The final ColNet model accurately segmented collagen fibrors, outperforming conventional intensity-based Otsu thresholding both qualitatively and quantitatively. ColNet achieved superior IoU and F1 scores compared to Otsu thresholding for all images (Fig. 2b,c and Fig. S7). Direct comparison of binary segmentation masks showed that ColNet successfully captures individual collagen fibrors that were missed by intensity thresholding, while avoiding false positive detection. A representative example of a softmax probability output layer of ColNet was provided as a confidence measure of the network (Fig 2d and Fig. S8). High softmax values (brighter) corresponded to well-defined fibror structures, while lower values (darker) indicated regions of uncertainty, typically associated with out-of-plane fibrors or areas with ambiguous signal. Additionally, ColNet performance was validated on a second unseen test dataset, generated by an independent annotator^45^, again outscoring Otsu intensity thresholding in IoU and F1 metrics (Fig. S9).

To demonstrate ColNet’s capability of volumetric collagen fibror segmentation, we deployed the model on a complete light-sheet dataset of a cleared desmoid tumor (Fig. 2e). ColNet segmentation was performed on each z-slice of the full dataset (311 z-slices, 2 µm step) and reconstructed as a 3D volume. High-magnification regions of interest revealed the model’s ability to accurately segment complex fibror architectures, as visualized through the softmax prediction layers. 3D surface reconstructions integrated the softmax predictions across the full tumor volume, illustrating the potential for global quantitative analysis of collagen network architecture.

### ColNet shows generalizability across diverse (fibrotic) tissues

A critical limitation of many deep learning models is their tendency to overfit to training data characteristics, learning dataset-specific characteristics rather than generalizable features. This overfitting restricts model applicability to new tissue contexts and typically requires extensive retraining. To assess whether ColNet learned generalizable features of collagen architecture rather than desmoid tumor-specific characteristics, ColNet was deployed on heterologous tissue samples that were not seen during training.

We first applied the modified DISCO clearing protocol to human skin biopsies from both healthy tissue and a healing wound at day 15 post-injury (Fig. 3). Samples were dual-stained with Fast Green FCF for collagen visualization and YO-PRO-1 for nucleus staining (Fig. S10). Similar to the desmoid tumor protocol, we limited Fast Green FCF staining duration to 24 hours to prevent oversaturation while maintaining adequate signal intensity, and employed continuous rotation during incubation to ensure uniform dye penetration. The clearing protocol enabled light-sheet imaging across a total z-depth of 850 μm, with samples imaged up to 6 mm deep in the dermis, clearly revealing the skin structure in three dimensions. YO-PRO-1 staining distinctly visualized the epidermis layer (Fig. 3a,d overview) and skin appendages present in the dermis (Fig. 3a,d insets), while Fast Green FCF highlighted the collagen-rich dermis underlying the epidermis (Fig. 3a,d overview). Notably, the collagen network was more densely packed underneath the re-epithelialized epidermis than at the wound edges, indicating active ECM remodelling in the wound^46^. Around skin appendages, collagen networks varied by structure: fibrors around a sweat gland were less organized and more dense (Fig. 3a), while a hair follicle was surrounded by concentrically organized collagen sheaths (Fig. 3d).

**Fig. 3.**
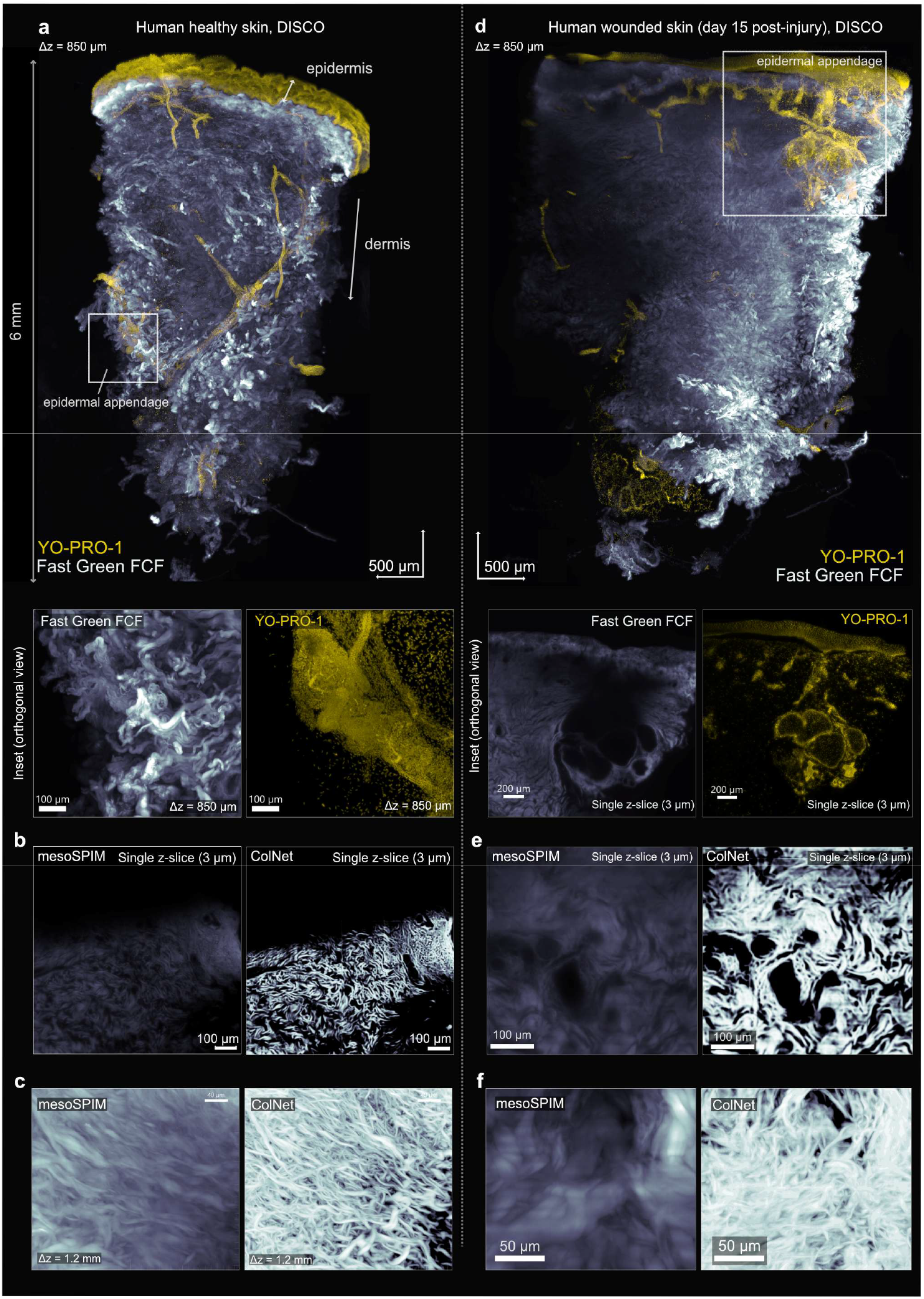
ColNet enables three-dimensional segmentation of collagen architecture in DISCO-cleared human skin biopsies. a, Light-sheet imaging (Benchtop mesoSPIM) of healthy human skin labeled with YO-PRO-1 (nuclei) and Fast Green FCF (collagen), showing epidermis with underlying dermis. Overview (perspective view) shows maximum intensity projection (MIP) spanning 850 µm z-depth (Δz = 850 µm); inset (orthogonal view) displays a sweat gland (MIP, 850 µm). b, c (different human healthy skin sample) ColNet accurately segments foreground collagen fibrors on single z-slices (b), resolving individual fibrors with improved signal-to-noise ratio in 3D (c). d, Light-sheet imaging (Benchtop mesoSPIM) of human skin 15 days post-injury, labeled with YO-PRO-1 (nuclei) and Fast Green FCF (collagen), showing epidermis with underlying dermis. Overview (perspective view) shows maximum intensity projection (MIP) spanning 850 µm z-depth; inset (orthogonal view) displays a sweat gland (single z-slice, 3 µm). Gamma correction (1.5 in Imaris) was applied on the Fast Green FCF channel for visualization. e,f ColNet accurately segments foreground collagen fibrors on single z-slices (e), resolving individual fibrors with improved signal-to-noise ratio in 3D (f).

Next, we assessed ColNet’s performance and generalizability on these skin samples. The model was directly applied on each z-slice (z=284 with 3 µm step) throughout the entire tissue volumes without any additional fine-tuning, retraining or hyperparameter adjustment (Fig. 3b,e). Segmented 2D slices were then reconstructed as a full z-stack (Fig. 3c,f). Side-by-side comparison of single z-slices between raw mesoSPIM input images and ColNet softmax output demonstrated effective segmentation of foreground collagen fibrors without oversegmenting out-of-focus structures (Fig. 3b,e). This segmentation strategy is crucial for accurate 3D reconstruction, as oversegmentation introduces axial blur when reconstructed in 3D (Fig. S11). Indeed, maximum intensity projections of ColNet softmax predictions revealed individual collagen fibrors with substantially improved signal-to-noise ratio compared to raw input images (Fig. 3c,f). Moreover, the resolved ColNet images recoveredfine fibror details that remained obscured by point spread function-induced blur in the original mesoSPIM acquisition images.

To further validate ColNet’s cross-tissue generalizability, we evaluated its performance on fibrotic mouse lung and liver tissue, derived from archived formalin-fixed paraffin-embedded (FFPE) material. We used fibrotic lungs (bleomycin-treated) and livers (CCl4-treated) from wild type mice. Samples were deparaffinized, washed in xylene and rehydrated in ethanol dilution series^47^ before applying the modified DISCO protocol with dual staining for collagen (Fast Green FCF) and nuclei (YO-PRO-1) (Fig. S12 and S13). We achieved successful clearing and imaging of both tissue types up to 2.6 mm depths (Fig. 4a,d). YO-PRO-1 delineated general tissue architecture, while Fast Green FCF revealed collagen organization across the entire tissue volume. In liver, low-magnification imaging of the YO-PRO-1 channel provided comprehensive visualization of liver structure (Fig. 4a), while higher magnification regions of interest displayed collagen signal in central and portal veins (Fig. 4b). ColNet deployment on a central vein clearly highlighted the collagen network architecture (Fig. 4c). In bleomycin-treated lungs, YO-PRO-1 delineated bronchi, while Fast Green FCF visualized both bronchial collagen deposition and pleural architecture (Fig. 4d). ColNet deployment again highlighted the collagen network within bronchi while averaging out signal intensity variations, providing clearer visualization of collagen organization (Fig. 4e).

**Fig. 4.**
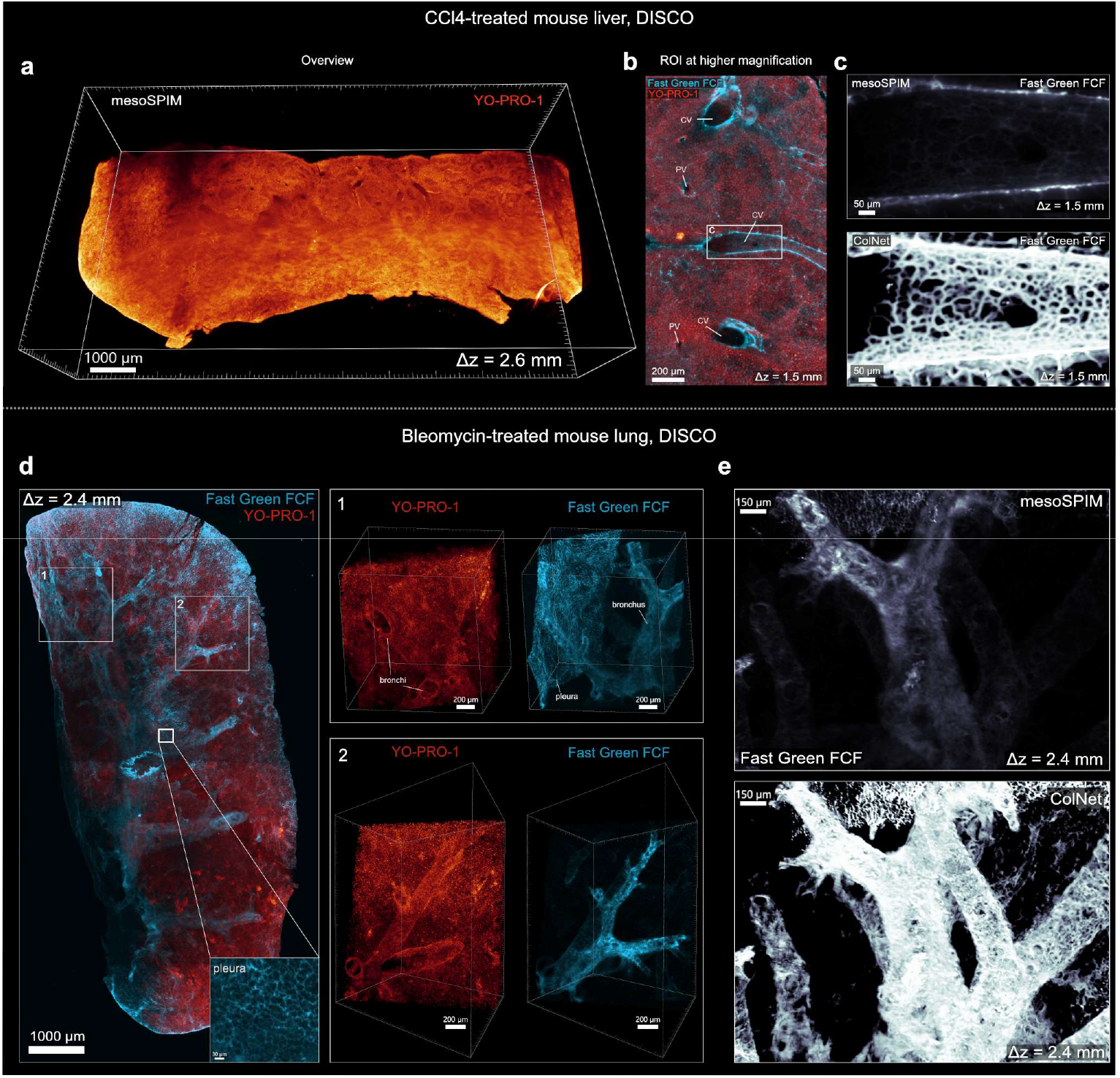
ColNet enables three-dimensional segmentation of collagen architecture in DISCO-cleared fibrotic mouse lung and liver tissue from archived FFPE material. a, Light-sheet imaging (Benchtop mesoSPIM) of CCl4-treated mouse liver derived from archived FFPE tissue, dual-labeled with YO-PRO-1 (nuclei) and Fast Green FCF (collagen). Overview shows maximum intensity projection (MIP) spanning 2.6 mm z-depth (Δz = 2.6 mm). b, Higher magnification region of interest (ROI) displaying MIP spanning 1.5 mm z-depth, revealing detailed collagen architecture within fibrotic regions. CV, central vein; PV: portal vein. c, MIP of ColNet softmax output demonstrating successful collagen fibror segmentation across the 1.5 mm volume. d, Light-sheet imaging (Benchtop mesoSPIM) of bleomycin-treated mouse lung derived from archived FFPE tissue, dual-labeled with YO-PRO-1 (nuclei) and Fast Green FCF (collagen). Two representative ROIs (inset 1 & 2) at higher magnification reveal bronchi and pleural structures across 2.4 mm z-depth. Gamma correction (1.5 in Imaris) was applied to the Fast Green FCF channel for visualization. e, 3D ColNet segmentation of collagen architecture in lung bronchi. Gamma correction (0.5 in Imaris) was applied to ColNet predictions for visualization.

Collectively, these results demonstrate that the modified DISCO protocol achieved effective clearing across diverse fibrotic tissue types. Moreover, ColNet maintained comparable segmentation performance across all tested tissues, confirming that the model learned general collagen architectural features, rather than desmoid-tumor specific characteristics from the original training dataset.

## DISCUSSION

Clearing collagen-rich tissues such as dermis, scar lesions, and desmoplastic tumors has remained a persistent bottleneck in optical tissue clearing for over a decade^23^. While methodological advances have been made^26,27^, no standardized pipeline currently exists for three-dimensional (3D) imaging and analysis of collagen architecture. Previous studies have focused either on the tissue clearing aspect or on computational analysis, but these approaches have not been integrated together. Here, using a *Xenopus* model of aggressive fibromatosis (desmoid tumors) as a proxy for collagen-rich tissues, we present a 3D pipeline combining optimized DISCO-based tissue clearing and mesoSPIM light-sheet fluorescence microscopy with deep learning-based feature extraction of collagen architecture.

### Optimized clearing of collagen-rich tissues

Building upon the methodology developed by Timin et al.^39^, we employ the Fast Green FCF and YO-PRO-1 dyes for collagen and nuclei labeling, respectively. We optimized clearing protocols for collagen-dense tissue using desmoid tumors as our technical platform. Key practicalfindings were that collagen-dense tissues require both high reagent volumes and extended dehydration times. These adaptations achieved optical transparency across *Xenopus* desmoid tumors, human skin biopsies, and fibrotic mouse lung and liver tissues. The Fast Green FCF and YO-PRO-1 combination proved particularly effective as a comprehensive visual readout: Fast Green FCF uniformly labels collagen architecture, while YO-PRO-1 delineates general tissue architecture.

Crucially, we established compatibility with archival formalin-fixed paraffin-embedded (FFPE) material through aDISCO-based processing^5^. Importantly, beyond the deparaffinization/xylene removal steps, no additional protocol modifications were required for FFPE samples, substantially broadening the method’s applicability to retrospective analysis of archived clinical material.

### Multimodal collagen imaging

Volumetric imaging on the Benchtop mesoSPIM platform enabled deep optical sectioning with penetration depths up to 2.6 mm in our cleared samples. Notably, we present here the first 3D high-resolution light-sheet recording of a desmoid tumor in its tumor microenvironment. Next, we validated compatibility of Fast Green FCF and YO-PRO-1 labelling with two-photon excitation using the Schmidt-Voigt objective^38^ in desmoid tumors. This represented the first deployment of the Schmidt-Voigt objective since its recent introduction and enabled high resolution collagen visualization through both fluorescence and second harmonic generation.

The Schmidt-Voigt objective resolvedfine collagen fibror organization, providing spatial detail that complemented the rapid volumetric acquisition capabilities of mesoSPIM. It’s multi-immersion design permits direct imaging of cleared samples in clearing media (dibenzylether in our case), eliminating any additional processing between light-sheet and two-photon acquisition. Importantly, while multi-photon excitation is compatible with both dyes, YO-PRO-1 exhibits SHG bleed-through, which should be considered when designing imaging experiments. This multimodal approach provides a versatile framework for investigating collagen architecture across multiple spatial scales and diverse tissue types.

### ColNet: a deep learning model for 3D collagen fibror segmentation

To address the lack of computational tools for 3D analysis of collagen organisation, we developed ColNet, a U-Net-based deep learning model for volumetric collagen fibror segmentation. The development of ColNet was facilitated by open-source platforms ZeroCostDL4Mic and DL4MicEverywhere which democratize deep learning for biomedical image segmentation through remote GPU connectivity, making deep learning accessible even to non-expert users^43,44^.

While recent work has demonstrated the value of U-Net architectures for collagen segmentation in second harmonic generation images of mouse skin^48^, achieving improved accuracy over intensity-based thresholding, the study was limited to 2D analysis of a single tissue type without 3D deployment or cross-tissue validation.

As intensity-based thresholding approaches cannot capture the fine structural details of collagen organization, we trained a 2D U-Net from scratch on volumetric light-sheet data from a cleared desmoid tumor. During model training, several critical technical observations emerged. First, ColNet trained optimally when provided with maximum contextual information, as largest patch sizes (within GPU memory constraints) showed the best convergence of training and validation losses. Second, optimal learning rate scheduling proved essential: rates below 5E-5 caused model instability and overfitting, resulting in oversegmentation, while ending training too early at higher learning rates led to undersegmentation. Third, qualitative performance assessment was crucial, as quantitative metrics alone can be misleading. When comparing learning rate scheduling, quantitative performance metrics were similar, yet qualitative performances were significantly different. Furthermore, despite a relatively low intersection-over-union (IoU) of 0.57, the softmax output exhibited high-quality segmentation of collagen architecture, demonstrating that traditional metrics may not always fully reflect model performance. These observations align with established principles of deep learning for biomedical image segmentation^21^.

An inherent challenge in this work was that collagen fibrors are difficult to annotate as ground truth. However, even with imperfect annotations, we employed a strategic approach to train the model on collagen fibror features using heterogeneous training images, allowing the model’s pa ern recognition capabilities to improve during training and effectively average out annotation imperfections.

Notably, ColNet demonstrated additional advantages beyond segmentation. Variations in illumination intensity across imaging depths are an inherent feature of light-sheet microscopy that can obscure structural features. While visualization tools such as DEVILS exist to correct these variations^49^, they modify pixel values non-linearly, precluding quantitative interpretation or feature extraction of processed images. In contrast, ColNet’s U-Net architecture averages out illumination differences across imaging depths during model training. This enables simultaneous visualization of structures obscured in raw images and feature extraction on the unmodified data. Furthermore, in human skin dermis, ColNet resolved fibrors with improved signal-to-noise ratio, highlighting structures that remained obscured by the point spread function in raw images. These results demonstrate that U-Net architectures can achieve robust performance with limited training data when combined with appropriate annotation strategies and optimized training parameters. This further validates previous observations^21^.

After obtaining The final model, ColNet was deployed across diverse biological contexts in 3D. Despite being trained exclusively on desmoid tumor data, ColNet maintained robust segmentation performance across human skin biopsies and fibrotic mouse lung & liver tissues. This cross-tissue generalizability is significant, demonstrating the model’s ability to recognize diverse collagen architectures without requiring retraining or hyperparameter fine-tuning.

### Conclusion and future perspectives

This work presents an integrated workflow combining tissue clearing, volumetric imaging, and deep learning for three-dimensional assessment of collagen architecture. ColNet’s cross-tissue segmentation accuracy demonstrates its potential for automated feature extraction of fibror orientation, alignment, density, and network topology. As such, our pipeline holds promise for quantitative investigations of cell-ECM dynamics that underlie many connective tissue diseases.

## MATERIALS & METHODS

### CRISPR/Cas9-mediated generation of desmoid tumors in *Xenopus*

All experiments involving *Xenopus* animals were conducted in accordance with local legal and institutional guidelines and approved by the governing ethical commi ee (EC2022-060). Ovulation was induced by injection of human chorionic gonadotropin (hCG) with priming doses of 10 units for males and 20 units for females, followed 24 hours later by boosting doses of 100 units for males and 150 units for females. Fertilized embryos were obtained by natural mating or in vitro fertilization. To generate mosaic apc mutants with desmoid tumors, *apc* sgRNA/Cas9 ribonucleoprotein complexes were microinjected into fertilized embryos as previously described^36^. Genotyping was performed using heteroduplex mobility assay and targeted amplicon deep sequencing as previously described^36,50^.

### Human skin samples

Sampling of cutaneous human wounds was approved by the Ethical Committee of Ghent University Hospital (EC2023/0106).

### Mouse FFPE samples

All experiments on mice were conducted in accordance with local legal and institutional guidelines and approved by the ethical committee of the Center for Inflammation Research and Ghent University, Faculty of Sciences.

### DISCO clearing and collagen staining of desmoid tumor samples

Following dissection, tissue samples werefixed in 4% paraformaldehyde (PFA) for 24 hours at room temperature. First dehydration was performed at room temperature in afive-step methanol (MeOH)/PBS series (20%, 40%, 60%, 80%, 100%; minimum 1 hour per step), followed by an additional overnight wash in 100% MeOH to ensure complete dehydration. Samples were bleached in 10% H2O2/MeOH solution at room temperature until depigmented (3-5 days), in a light box on a rocking table at room temperature. Following bleaching, samples were rehydrated at room temperature through a reversefive-step graded PBS/MeOH series. Samples were then permeabilized with three 1-hour washes in 0.5% PBS-Triton-X-100 (PBS-T) at room temperature and one overnight wash at 4°C, all performed with continuous rocking. Nuclear staining was achieved by incubating samples with YO-PRO-1 (3:1000 dilution) (Invitrogen, Cat#Y3603) in 0.2% PBS-T for 24 hours at room temperature on a rotating wheel. To make the mounting easier, samples can optionally be embedded in 2% low-melting point agarose in MilliQ water and washed for 1 hour in 1x PBS at room temperature on a rocking table. A second dehydration series was performed at room temperature using a 5-step (20%, 40%, 60%, 80%, 100%; minimum 1 hour per step) Milli-Q water/MeOH solutions. Upon reaching 100% MeOH, samples underwent two additional 1-hour 100% MeOH washes followed by an overnight 100 % wash at room temperature with rocking to ensure complete dehydration. Collagen staining was performed with Fast Green FCF (2 µg/mL) (Thermo Scientific, Cat#A16520.22) in 100% MeOH for 24 hours at room temperature on a rotating wheel, followed by a 1-hour wash in 100% MeOH after staining^39^. Samples were then treated with 100% dichloromethane (DCM) for two 1-hour incubations at room temperature with shaking, followed by an overnight incubation in 100% DCM at room temperature. Finally, samples were cleared in dibenzyl ether (DBE) with two initial 1-hour washes, then stored in fresh DBE until imaging.

### DISCO clearing and collagen staining of human skin samples

Following full-thickness skin biopsy, tissue samples werefixed in 4% paraformaldehyde (PFA) for 24 hours at 4°C. Thefirst dehydration was performed at 4°C in afive-step MeOH)/PBS-Triton-X-100-Glycine (PBS-TG: 0.1% Triton-X-100, 0.3 M glycine in 1x PBS) series (20%, 40%, 60%, 80%, 100% MeOH; minimum 1 hour per step), followed by two 1-hour washes in 100% MeOH and an additional overnight wash at 4°C in 100% MeOH to ensure complete dehydration. Thefirst delipidation was performed by incubating samples in 100% DCM for two 1-hour washes at room temperature with shaking, followed by an additional overnight incubation in 100% DCM at room temperature. Samples were then transferred to 100% MeOH for two 1-hour washes at 4°C. Bleaching was performed at room temperature in 5% H2O2/MeOH solution until samples were depigmented (3–5 days), in a light box on a rocking table. Following bleaching, samples were rehydrated at 4°C through a reverse five-step graded PBS-TG/MeOH series (80%, 60%, 40%, 20% MeOH, then 100% PBS-TG; minimum

1 hour per step). Before permeabilization, samples can be washed with 0.1M KOH in 0.5% PBS-T, to enhance reagent penetration. Samples were then permeabilized with three 1-hour washes in PBS-TG at room temperature and one overnight wash at 4°C, all performed with continuous rocking. An additional 1-hour wash was performed at room temperature in PTxWH buffer (PBS containing 0.1% Triton-X-100, 0.05% Tween-20, and 2 µg/mL heparin). Nuclear staining was achieved by incubating samples with YO-PRO-1 (3:1000 dilution; Thermo Fisher Scientific) in 0.2% PBS-T for 24 hours at room temperature on a rotating wheel. Following nuclear staining, samples were embedded in 2% low-melting point agarose prepared in Milli-Q water and washed for 1 hour in 1x PBS at room temperature on a rocking table. A second dehydration series was performed at room temperature using a graded Milli-Q water/MeOH series (20%, 40%, 60%, 80%, 100% MeOH; minimum 1 hour per step). Upon reaching 100% MeOH, samples underwent two additional 1-hour washes in 100% MeOH followed by an overnight wash at room temperature with rocking to ensure complete dehydration. Collagen staining was performed by incubating samples with Fast Green FCF (2 µg/mL) in 100% MeOH for 24 hours at room temperature on a rotating wheel, followed by a 1-hour wash in 100% MeOH (Timin & Milinkovitch, 2023). The second delipidation was performed by treating samples with 100% DCM for two 1-hour incubations at room temperature with shaking, followed by an overnight incubation in 100% DCM at room temperature. Finally, samples were cleared in DBE with two initial 1-hour washes, then stored in fresh DBE at room temperature until imaging.

### DISCO clearing and collagen staining of archived FFPE samples

Following dissection, tissue samples werefixed in 4% paraformaldehyde (PFA) for 24 hours at room temperature. For formalin-fixed paraffin-embedded (FFPE) samples, deparaffinization and rehydration were performed as described in the aDISCO paper^51^. Tissue blocks were first melted for 1 hour at 60°C. Samples were incubated in xylene to remove remaining wax (two 1-hour incubations at 37°C with shaking), then rehydrated through a seven-step graded ethanol (EtOH)/Milli-Q water series (100%, 95%, 90%, 80%, 70%, 50%, 25%; minimum 1 hour per step), followed by an overnight incubation in 1x phosphate-buffered saline at room temperature while shaking. All samples were then washed for 1 hour in 1x PBS. The remaining steps were performed as described for desmoid tumors (cf. *‘DISCO clearing and collagen staining of desmoid tumor samples’*).

### Microscopy and imaging

For light-sheet imaging, cleared samples (desmoid tumors, lungs, liver, skin) were imaged using selective plane illumination microscopy (SPIM) on the Benchtop mesoSPIM platform in axially swept light-sheet microscopy (ASLM) mode for uniform axial resolution across thefield of view. The excitation path utilized an Oxxius L4Cc laser combiner with 405, 488, 561, and 638 nm laser lines. YO-PRO-1 (Invitrogen, Cat#Y3603) nuclear staining and Fast Green FCF (Thermo Scientific, Cat#A16520.22) collagen staining were excited using the 488 and 638 nm laser lines, respectively. The detection path was equipped with Mitutoyo Plan Apo BD objectives (2x/0.10, WD 34 mm; 5x/0.14, WD 34 mm; 10x/0.28, WD 33.5 mm) paired with a Mitutoyo MT-1 tube lens (f = 200 mm). Fluorescence emission was collected through a Teledyne Photometrics Iris 15 sCMOS camera (sensor diagonal 25 mm, pixel size 4.25 µm, 15 megapixels). Camera exposure time was 20 ms. Default galvo scanning seetings (99.9 Hz) were used for all mesoSPIM recordings. For two-photon imaging (desmoid tumors), the Schmidt-Voigt objective was used as described in the original paper^38^. A femtosecond Ti:sapphire laser (Chameleon Ultra II, Coherent) tuned to 925 nm provided excitation for second harmonic generation (SHG) imaging and collagen visualization. SHG signals were detected using a 470/40 nm bandpassfilter. Fast Green FCF and YO-PRO-1 staining were excited at 940 nm and 900 nm, respectively, with a 470/40 bandpassfilter. For both imaging systems (light-sheet & two-photon), samples were embedded in 2% low-melting point agarose and mounted in custom 3D-printed sample holders. All cleared samples were immersed in dibenzyl ether (DBE, RI = 1.56) as the refractive index matching medium during imaging. Detailed acquisition parameters (objective, laser power, z planes z-step size …) for each sample are provided in Supplementary Table 1. For multi-tile acquisitions, stitching was performed with BigStitcher^52^. Data were rendered with Fiji^53^ or Imaris (Oxford Instruments).

### Deep learning

The ColNet deep learning model was trained from scratch to segment collagen fibrors using the ZeroCostDL4Mic platform^43^. The model was based on the classical 2D-U-Net architecture^32^ and trained on ten manually annotated 1536×1536 images (n=10) from a Fast Green FCF-stained DISCO-cleared desmoid tumor dataset, with corresponding binary masks (8-bit) created in Fiji. The patch size of 1536×1536 pixels was selected as the maximum size that could be processed given available GPU memory constraints of 24 GB vRAM; larger patch sizes also provided more contextual information and improved training stability without overfitting. To correct for selection bias, a Fiji macro was used to randomly extract 1536×1536 patches from z-stacks across the entire sample volume (z=311). Of these, 8 images were used for training and 2 for validation during training. The standard U-Net architecture was used^32^. Dropout (0.5) was applied after the deepest convolutional layers. The final layer used a 1×1 convolution with sigmoid activation to produce binary segmentation masks. The network contained a total of 31,031,685 trainable parameters. Training was performed on an NVIDIA GeForce RTX 4090 GPU (24 GB vRAM) using TensorFlow (v2.12.0), Keras (v2.12.0), NumPy (v1.22.4), and CUDA (v11.8.89). The model was trained with a weighted binary cross-entropy loss function, with class weights calculated inversely proportional to class frequencies in the training dataset. Training parameters included a batch size of 1 and 9 steps per epoch. Data augmentation was applied during training, including random rotations, horizontal and vertical flipping, zoom magnification, shifting, and image shearing. The final training protocol (Model 3) consisted of 1,050 epochs (9,450 iterations) with a multi-step learning rate schedule using the Adam optimizer: 500 epochs (4,500 iterations) at 1E-4, 250 epochs (2,250 iterations) at 5E-5, 100 epochs (900 iterations) at 2E-5, and 200 epochs (1,800 iterations) at 1E-5. Additional training hyperparameters included a minimum fraction of 0.01 for patch acceptance. For quality control, nine additional images with variable patch sizes were manually annotated with corresponding masks. Model performance on this validation dataset achieved afinal Intersection over Union (IoU) of 0.57 and Dice coefficient of 0.72. For visualization and quantitative analysis, softmax layers were thresholded empirically to optimize segmentation quality and minimize false positives. Upon deployment of ColNet to new datasets, no fine-tuning was performed. Prior to prediction, a small blob (4×4 pixels) with the median pixel intensity of the entire volume was optionally applied to each z-slice to stabilize normalization across the dataset.

## Contributions

W.H., K.V. and T.N. conceptualized the study and designed the experiments. W.H. conducted the majority of the experimental work. L.S. and N.B. performed the experimental work on skin samples. M.B. performed CRISPR design and microinjections for the desmoid tumor *Xenopus* model. L.S. performed mesoSPIM imaging of human skin samples in Fig. S7 & S8. T.N. performed all other mesoSPIM imaging. W.H. and T.N. performed image analysis. A.D.V. conducted all Schmidt-Voigt microscopy. A.D.V., N.V., F.F.V. and F.H. provided expertise and support for mesoSPIM and Schmidt-Voigt microscopy. L.S., M.G., P.B. and E.H. provided skin samples. B.V. and C.L.S. provided archived FFPE mouse liver samples. C.M.B.G. and B.N.L. provided archived FFPE mouse lung samples. W.H. wrote the original draft with input from T.N. and K.V. T.N. and K.V. reviewed and edited the manuscript. T.N. and K.V. led project supervision.

## Acknowledgment

T.N. received funding from the ‘Bijzonder Onderzoeksfonds’ of Ghent University (01P06323), the Research Foundation Flanders (1294725N) and was further supported by a ‘young investigator proof-of-concept’ (YIPOC) grant from the Cancer Research Institute Ghent (CRIG). Research in the K.V. laboratory is supported by the Research Foundation— Flanders (FWO-Vlaanderen) (3G0A6922), by the Foundation against Cancer (365L07823) and by the Concerted Research Actions from Ghent University (01G02320). Further support was obtained by the Desmoid Tumor Research Foundation, the Desmoid Tumor Foundation of Canada and SOS Desmoïde. Research in the E.H. lab is funded by an FWO grant (3G071224), by Ghent University Research Fund (BOF/24J/2023/138), by the LEO Foundation (SE-24-800055) and by Foundation against Cancer (365L04523). F.F.V. is funded by a Branco Weiss Fellowship, Society in Science, administered by the ETH Zurich. This work was further supported by the University Research Priority Program (URPP) “Adaptive Brain Circuits in Development and Learning (AdaBD)” of the University of Zurich (N.V., A.D.V. and F.H.).

## SUPPLEMENTARY FIGURES

**Fig. S1.**
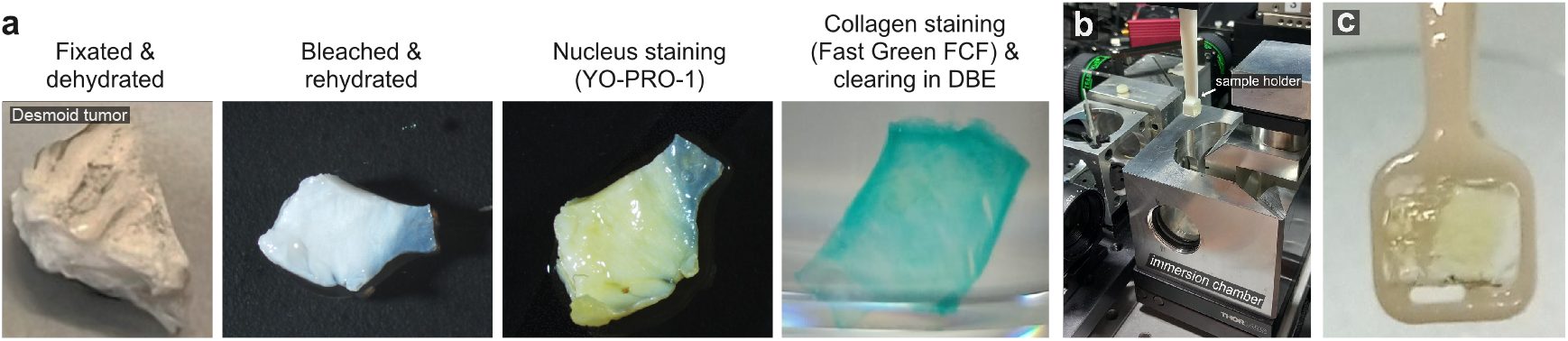
DISCO-cleared and stained desmoid tumor and Schmidt-Voigt objective setup. **a**, Macroscopic images of the tissue clearing process applied on desmoid tumors, with collagen staining (Fast Green FCF) and nucleus staining (YO-PRO-1). **b**, Schmidt-Voigt objective setup for collagen imaging using two-photon microscopy, **c**, Sample holder with mounted cleared desmoid tumor sample.

**Fig. S2.**
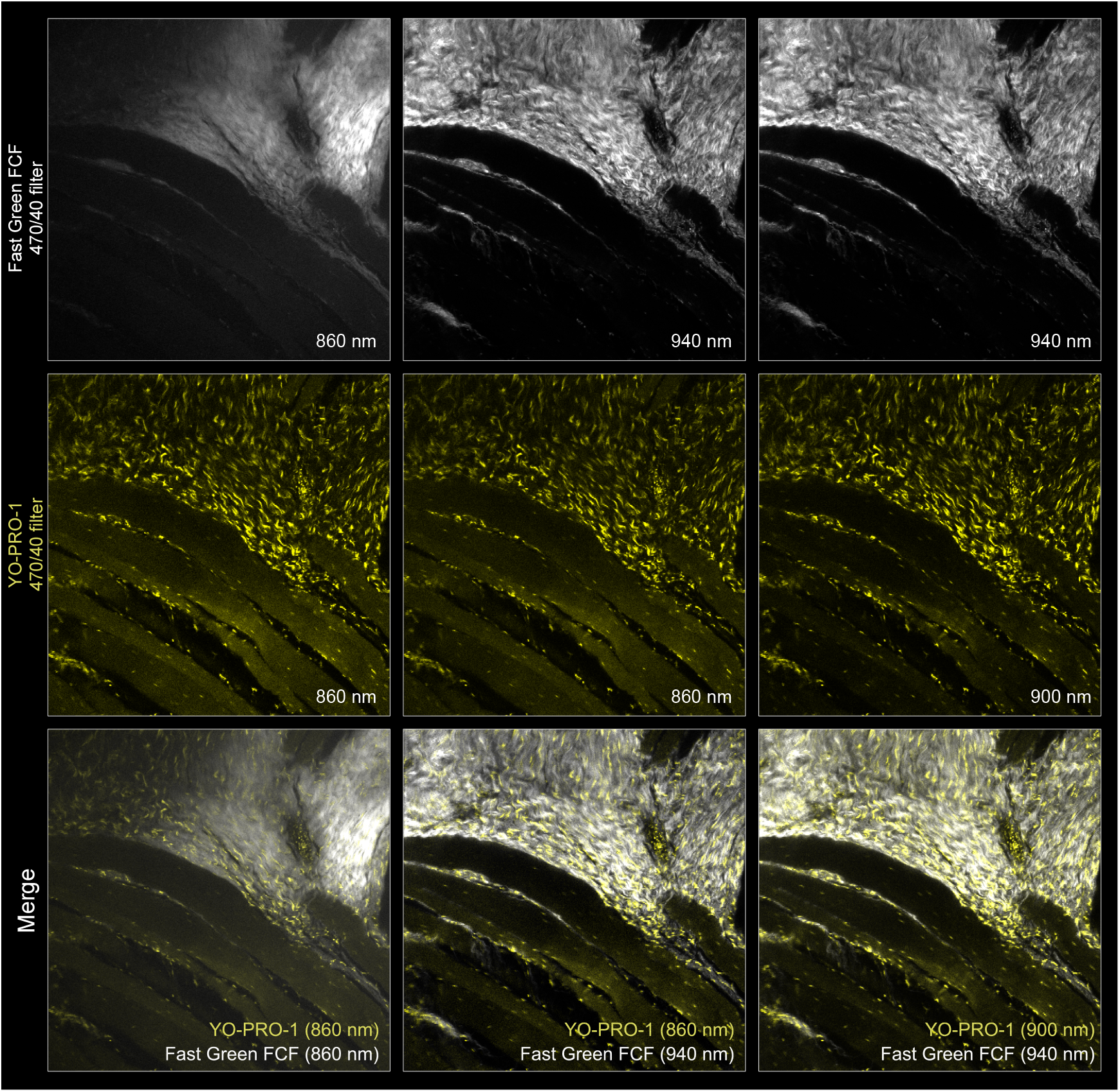
Two-photon excitation wavelength optimization for Fast Green FCF and YO-PRO-1. Systematic comparison of fluorescence intensity and image quality across different excitation wavelengths for collagen (Fast Green FCF) and nuclear (YO-PRO-1) visualization in DISCO-cleared *Xenopus* desmoid tumor tissue. Excitation at 940 nm (Fast Green FCF) and 900 nm (YO-PRO-1) provided opmaltimal signal-to-noise ratios. However, YO-PRO-1 multi-photon excitation showed spectral bleedthrough of the SHG signal, which should be considered when interpreing nuclear channel data in collagen-rich regions. Both channels were acquired on 470/40 filter.

**Fig. S3.**
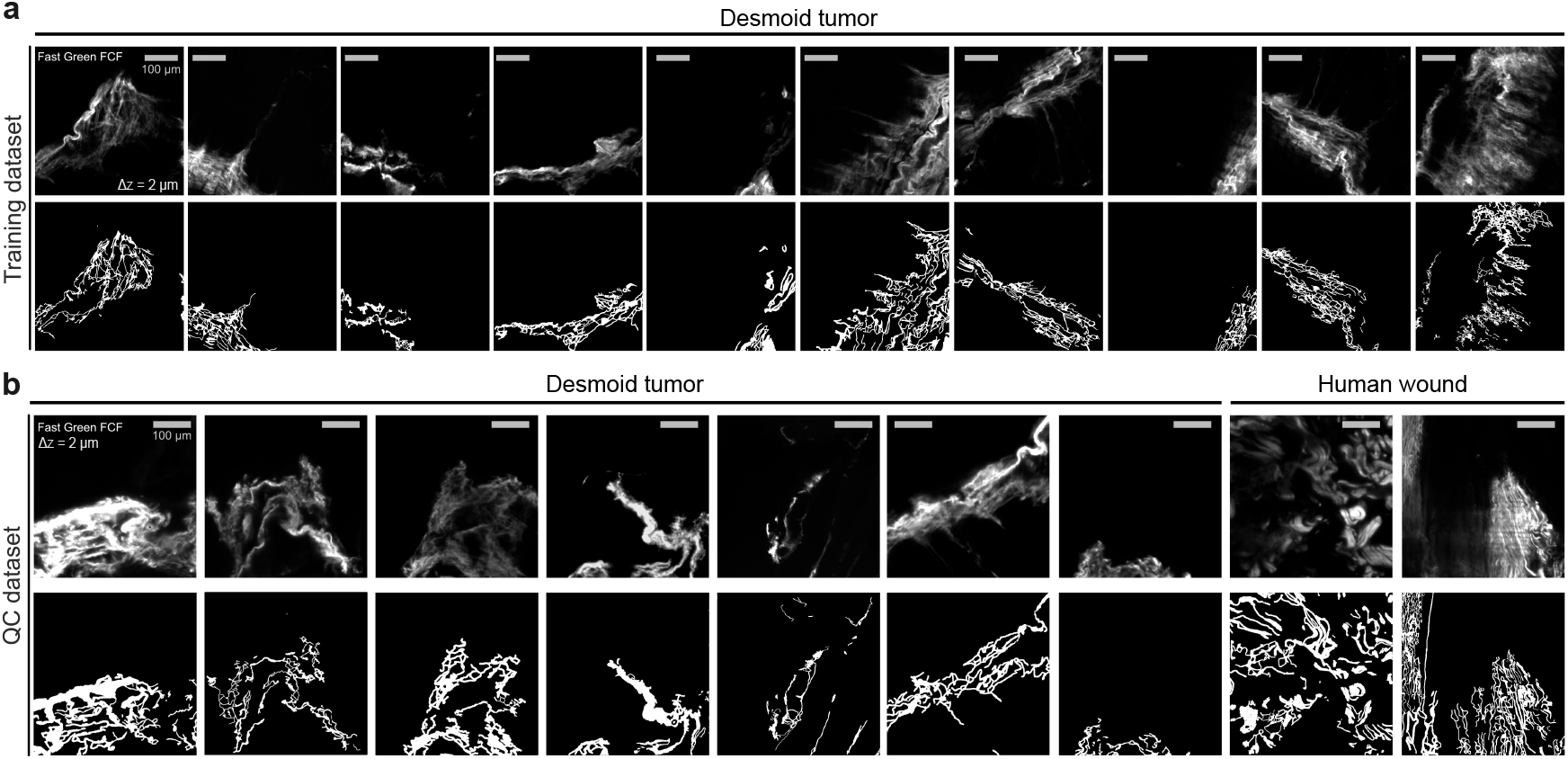
Training and quality control datasets with corresponding ground truth annotations for ColNet development and validation. **a**, Training dataset used for training of ColNet. Ten 1536×1536 pixel patches were randomly sampled from a light-sheet microscopy recording of a *Xenopus* desmoid tumor (311 z-planes spanning 622 µm) for model training. **b**, Quality control dataset used for validation of ColNet. An additional nine images with variable patch sizes were randomly sampled from the same volume for quality control assessment. To evaluate model generalizability, images from a human full thickness wound were included in the quality control dataset. Collagen structures were manually annotated in Fiji and exported as binary segmentation masks (8-bit) to serve as ground truth for supervised model training and validation.

**Fig. S4.**
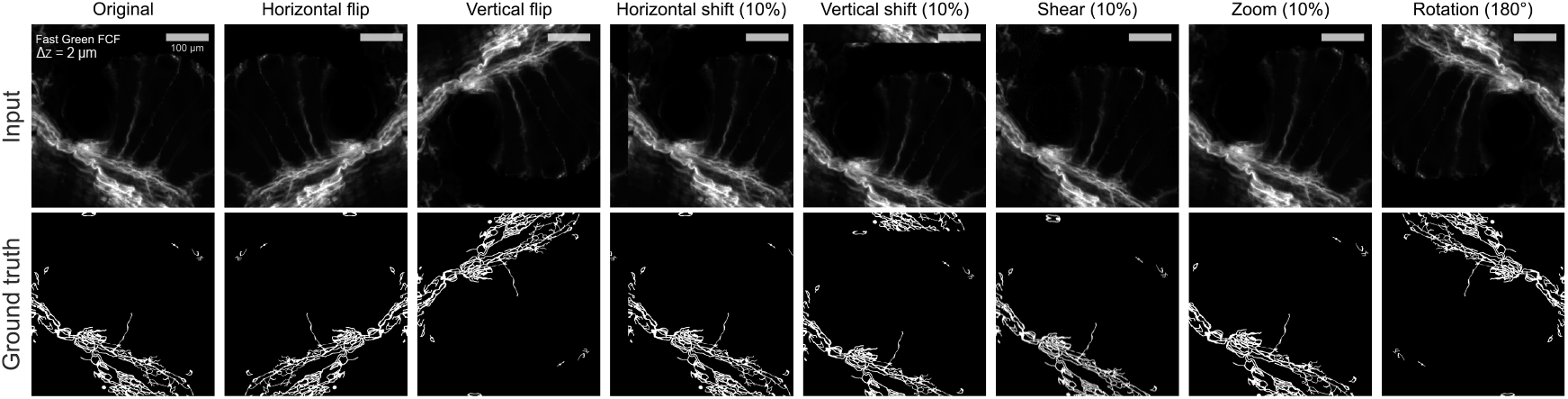
Data augmentation used during training of ColNet. Representiave example of augmentation transformaons applied to a single training image and its ground truth masks (8-bit) to increase model robustness and account for natural biological variance. Augmentations include: horizontal shi (10%), vercal shi (10%), horizontal flip (50% probability), vertical flip (50% probability), shear transformation (10%), zoom (10%), and rotation (0–180°).

**Fig. S5.**
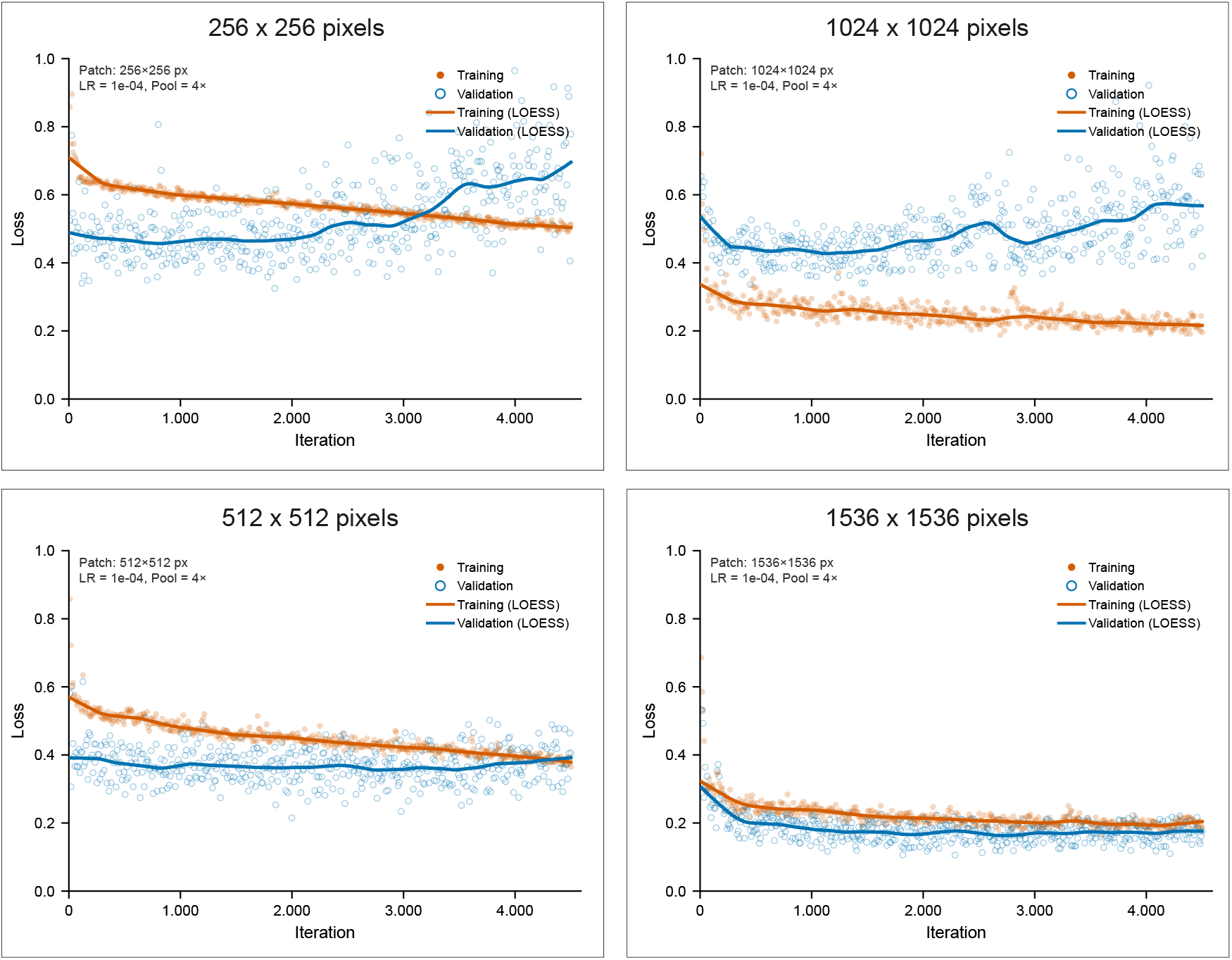
Effect of patch size on ColNet training dynamics. Two independent optimization experiments, each trained for 500 epochs at constant learning rate (1×10−4) with identical data augmentation as the final model (rotation 180°, horizontal and vertical flipping 10%, zoom 10%, horizontal and vercal shiftiing 10%, shearing 10%). Evaluated at batch size 1 with 4 max-pooling steps. Training and validation loss curves with LOESS smoothing for 256×256, 512×512, 1024×1024 and 1536×1536 pixel patches. Loss values above 1 were clamped for visualization. Small patch sizes exhibit clear overfing with divergent training and validation losses, while larger patches show progressively tighter convergence between training and validation, indicaing improved generalization through increased contextual information.

**Fig. S6.**
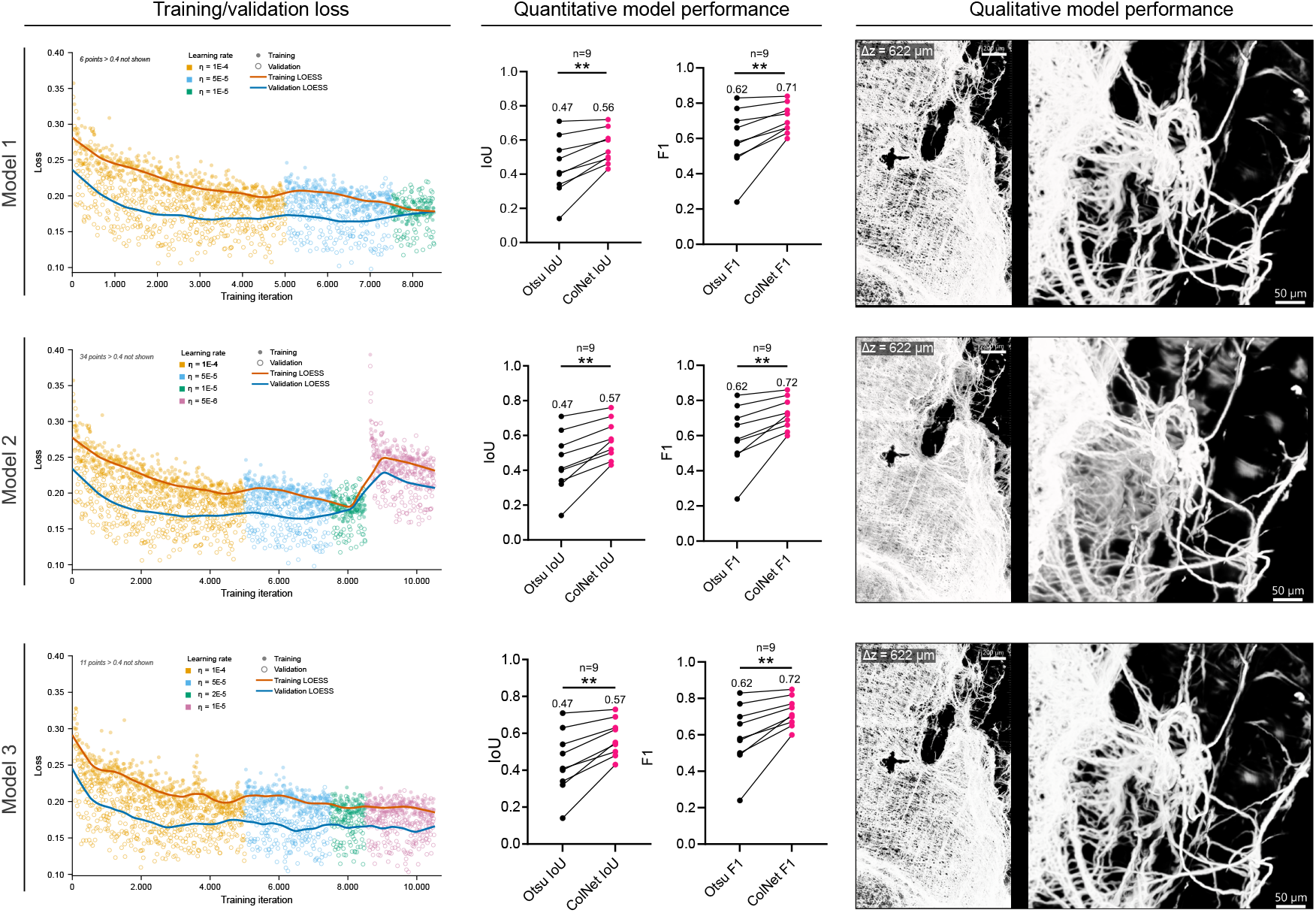
Impact of learning rate scheduling on ColNet training dynamics and segmentation quality. Three schedules tested (iterations/learning rate): Model 1 (4500/1E-4, 2250/5E-5, 900/1E-5); Model 2 (4500/1E-4, 2250/5E-5, 900/1E-5, 1800/5E-6); Model 3 (4500/1E-4, 2250/5E-5, 900/2E-5, 1800/1E-5). Le: Training and validation loss with LOESS smoothing, color-coded by learning rate. Loss values above 0.4 were clamped for visualization. Model 1 converges early at 1E-5, Model 2 exhibits training instability at ultra-low learning rate (5E-6), and Model 3 demonstrates stable convergence. Center: Test dataset IoU and Dice scores are similar across all three models, demonstraing that quantative overlap metrics alone can be insufficient for model selection (t-test, p<0.05, n=9). Right: 3D segmentation predictions with high-magnification ROI (inset) reveal substantial qualitative differences. Model 1 undersegments due to early convergence, Model 2 oversegments with false positives following unstable training, while Model 3 provides optimal segmentation with accurate fiber detection and specificity. Model 3 was selected for downstream analyses.

**Fig. S7.**
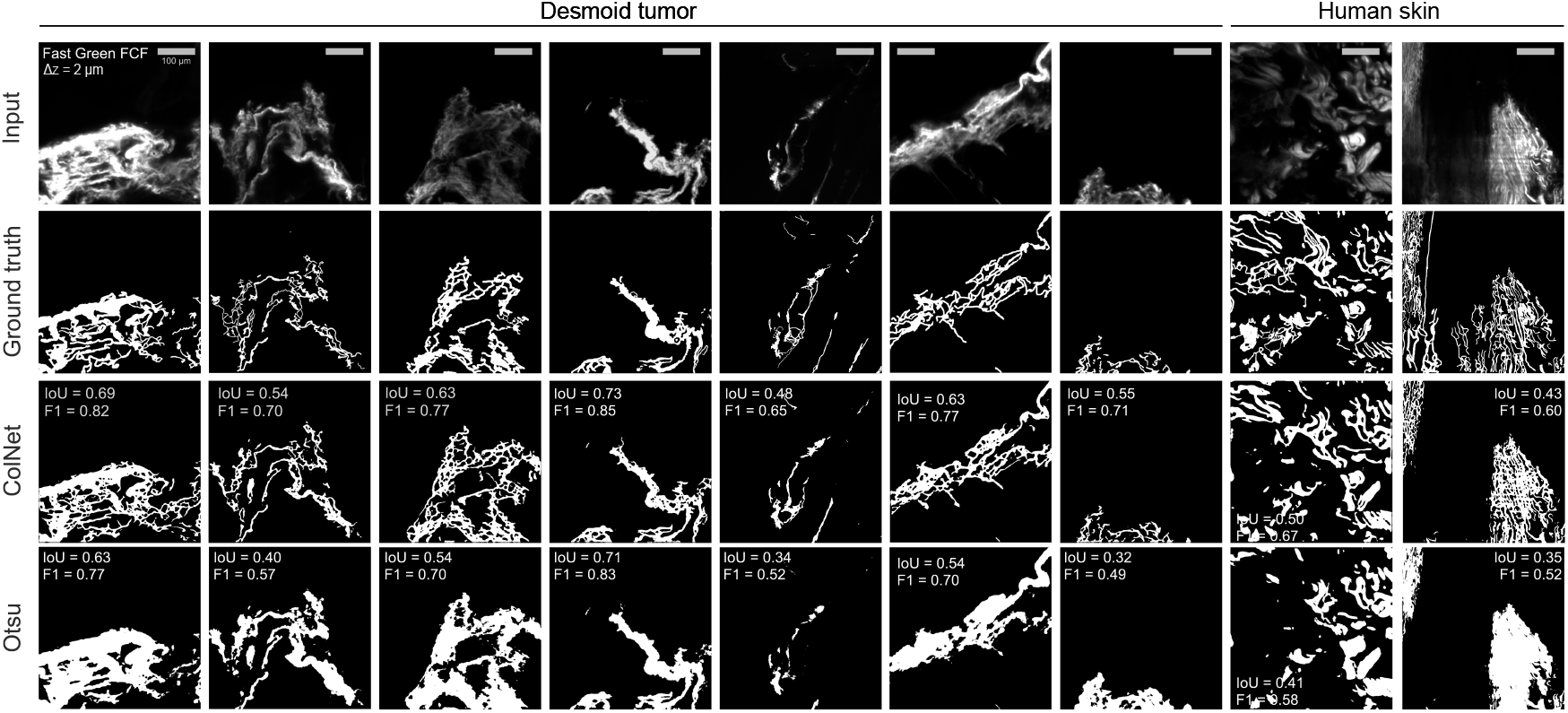
ColNet segmentation outperforms Otsu intensity thresholding across the complete unseen quality control dataset. Comparison of Otsu intensity thresholding versus ColNet segmentation on an unseen quality control dataset. For each segmentation mask (8-bit), Intersection Over Union (IoU) and F1 (Dice) score are displayed, demonstrating superior ColNet performance across diverse imaging conditions and tissue sources.

**Fig. S8.**
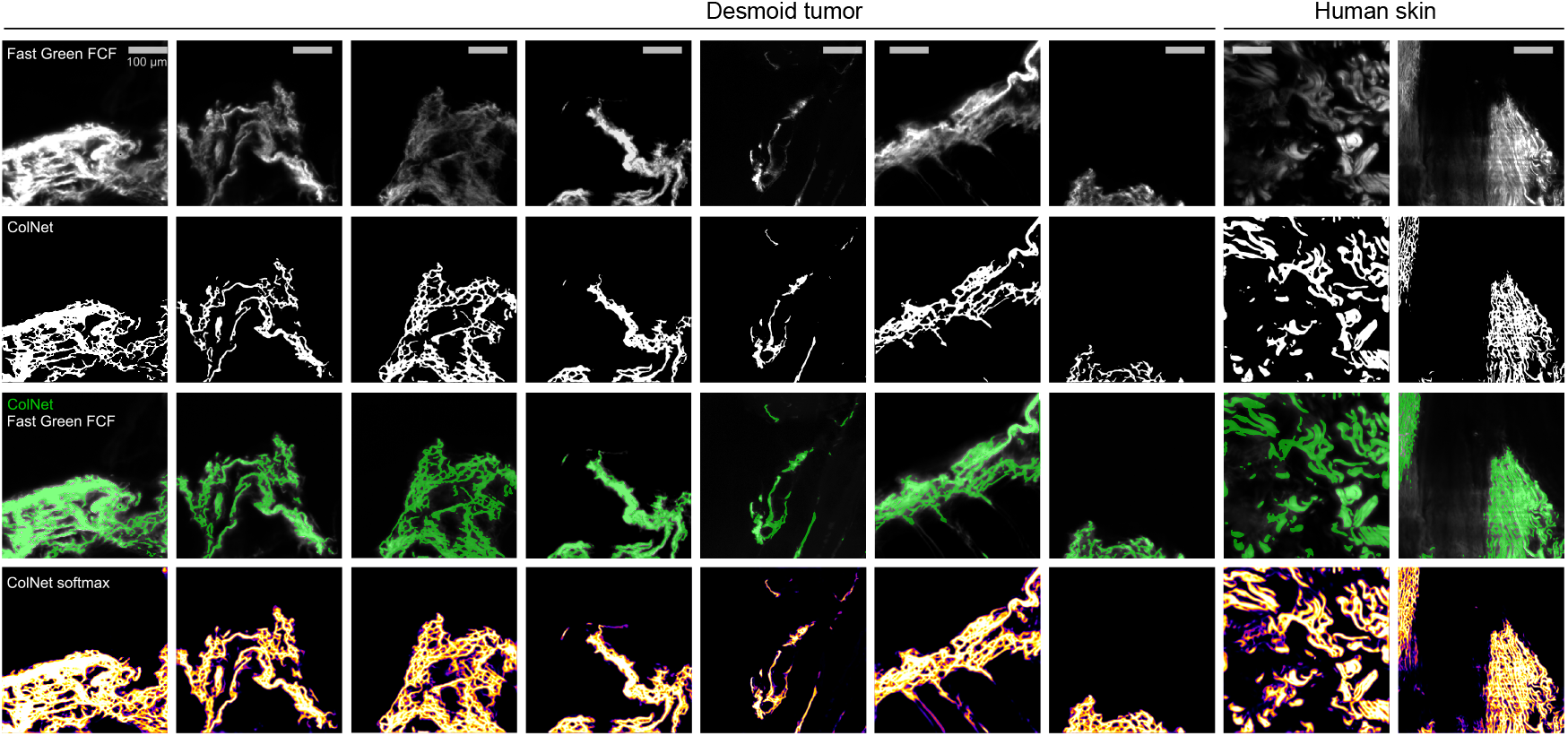
ColNet probabilistic output and segmentation performance on the complete unseen quality control dataset. Representative input image with corresponding binary segmentation masks (optimal threshold), overlay, and raw softmax probability layer demonstrating pixel-wise classification confidence. The softmax acvtiation funcon outputs a continuous probability distribution (0–1 scale) for each pixel. The original 16-bit grayscale somax map was pseudocolored with Fiji ‘Fire’ LUT for visualization, where warmer colors indicate higher model certainty for positive collagen class prediction.

**Fig. S9.**
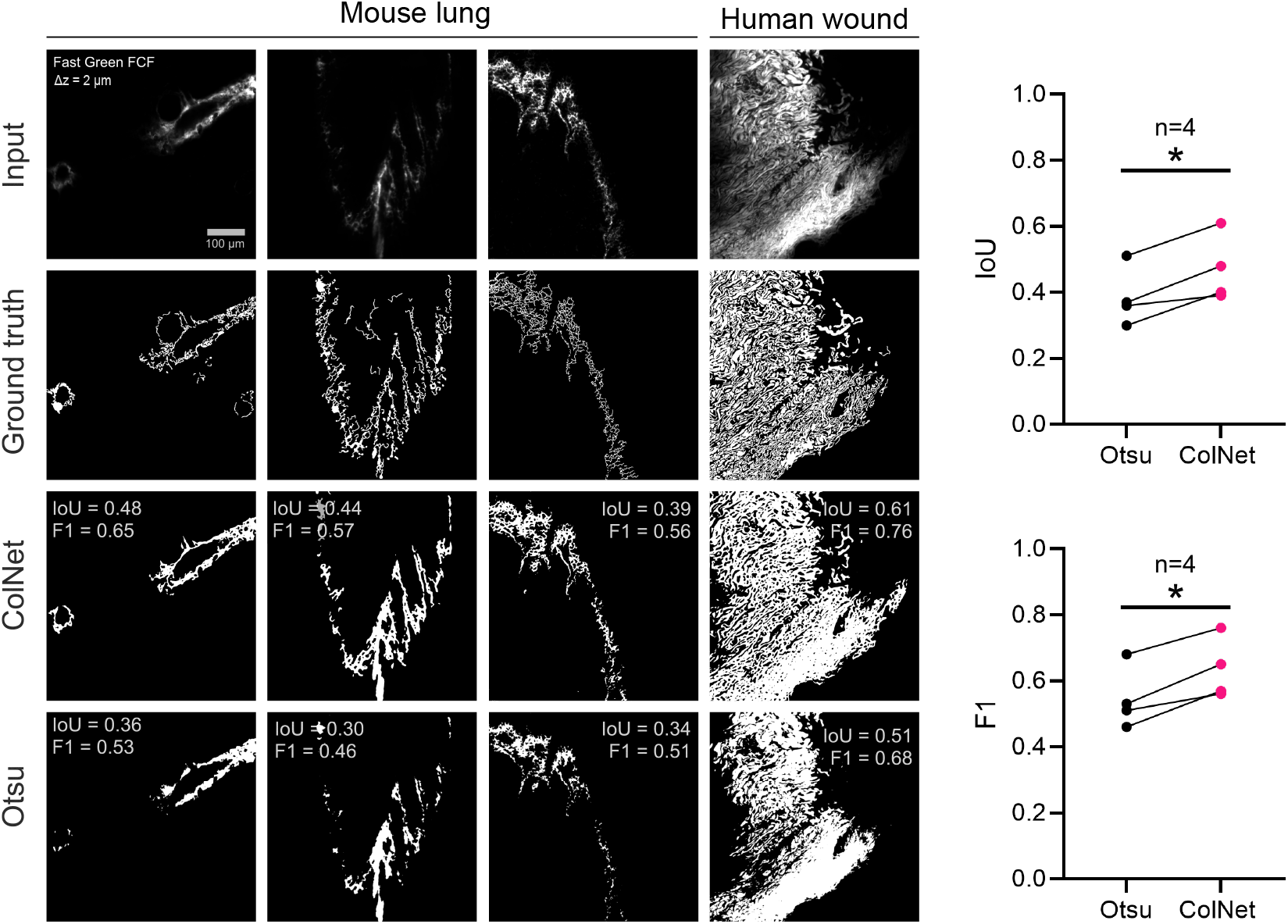
ColNet segmentation outperforms Otsu intensity thresholding across an independently annotated quality control dataset. Comparison of Otsu intensity thresholding versus ColNet segmentation on an independent, unseen quality control dataset. For each segmentation mask (8-bit), Intersection Over Union (IoU) and F1 (Dice) score are displayed, demonstrating superior ColNet performance across diverse imaging condions and ssue sources.

**Fig. S10.**
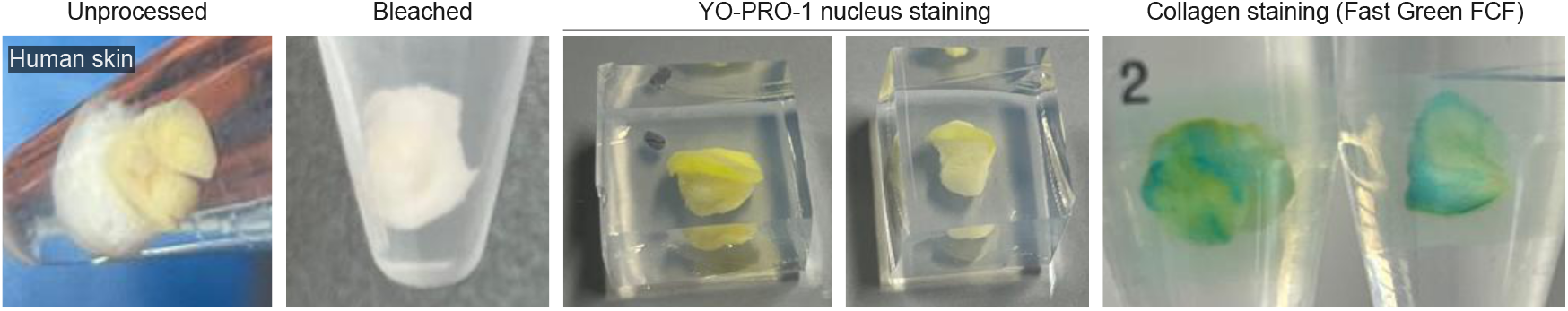
DISCO-clearing and staining of human skin. Representative macroscopic images of the tissue clearing process, applied on human skin bopsies, with collagen staining (Fast Green FCF) and nuclei (YO-PRO-1).

**Fig. S11.**
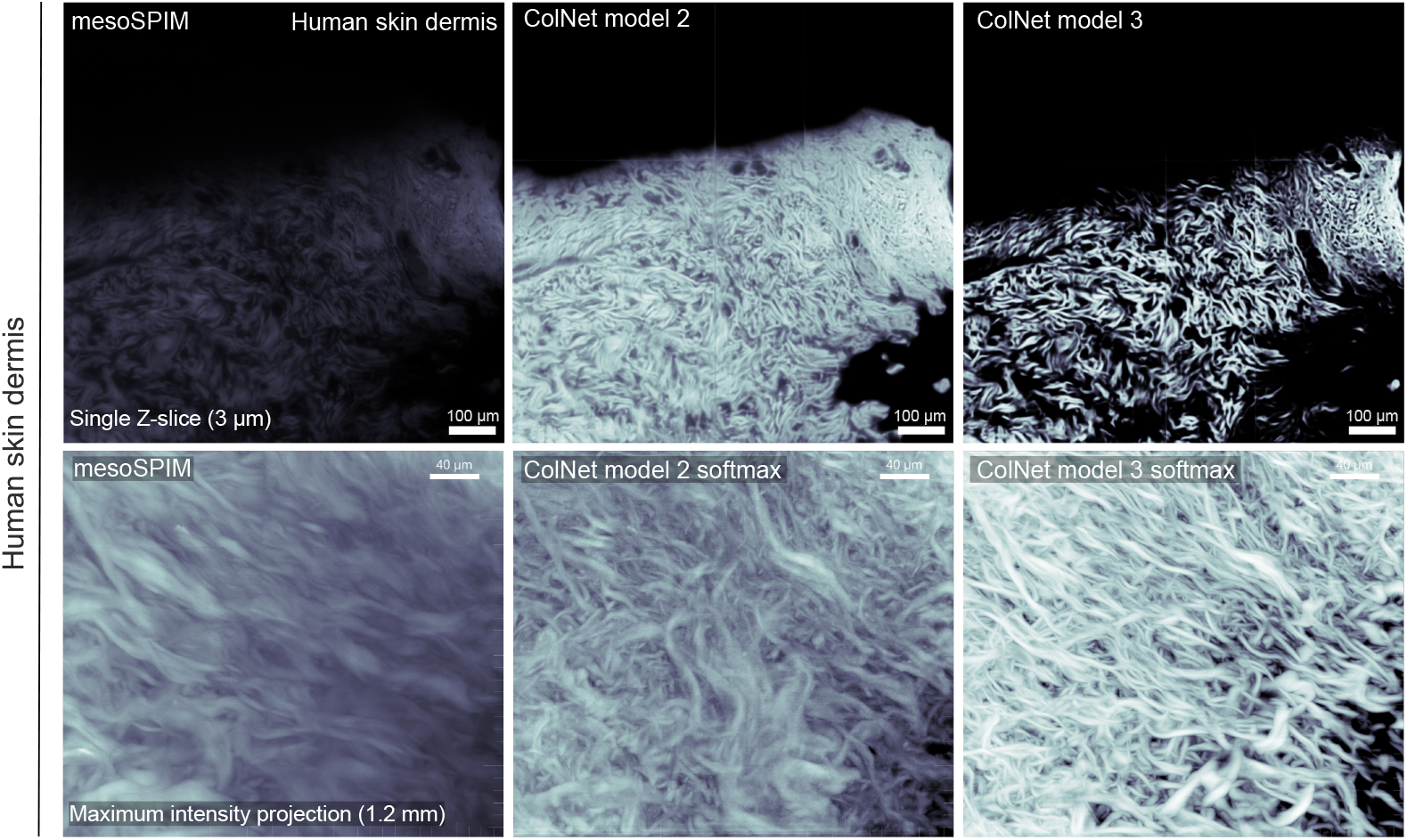
Effect of ColNet segmentation strategy on 3D softmax prediction in human skin dermis. Comparison of early and final ColNet softmax prediction layers on single z-slices and maximum intensity projections. The early model (Model 2) oversegmented individual z-slices, producing axial blur artifacts in 3D projections. The final model (Model 3) selectively segments foreground collagen fibers, yielding cleaner 3D projections with improved signal-to-noise ratio and enhanced resolution of individual fiber structures.

**Fig. S12.**
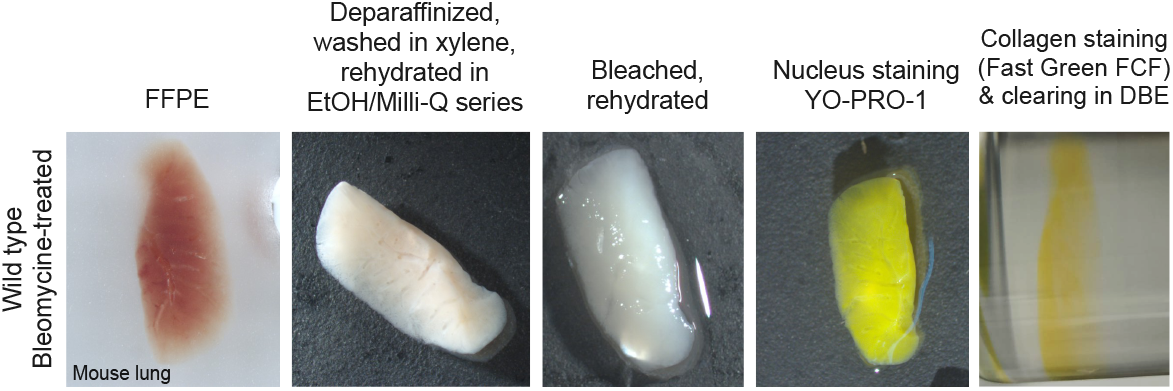
DISCO clearing and staining of archived FFPE mouse lung tissue. Representative macroscopic images of the tissue clearing process applied on mouse lungs, with collagen staining (Fast Green FCF) and nuclei (YO-PRO-1). FFPE: formalin-fixed paraffin-embedded.

**Fig. S13.**
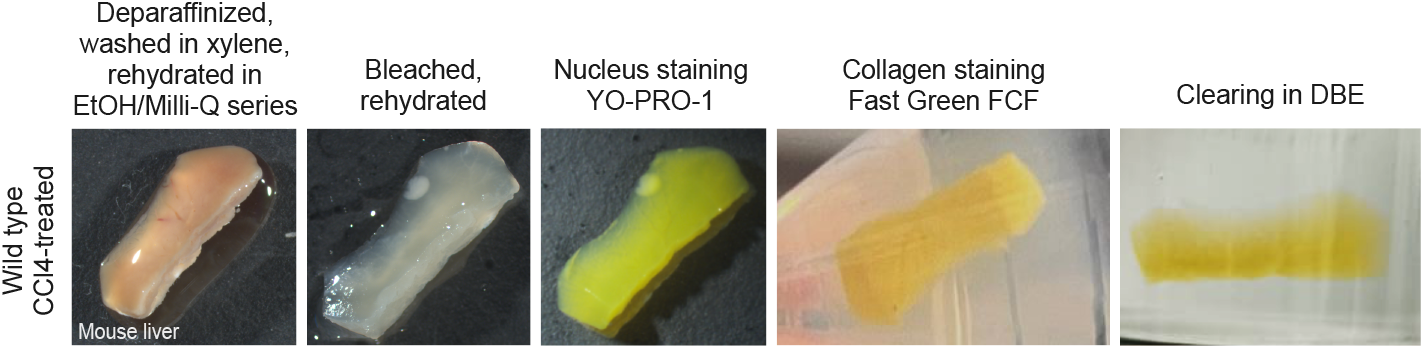
DISCO clearing and staining of archived FFPE mouse liver tissue. Representative macroscopic images of the tissue clearing process applied on mouse liver, with collagen staining (Fast Green FCF) and nuclei (YO-PRO-1).

